# A Third COVID-19 Vaccine Dose in Kidney Transplant Recipients Induces Antibody Response to Vaccine and Omicron Variants but Shows Limited Ig Subclass Switching

**DOI:** 10.1101/2024.09.01.610689

**Authors:** Jenny M. Lee, Jaiprasath Sachithanandham, John S. Lee, Janna R. Shapiro, Maggie Li, Ioannis Sitaris, Stephanie R. Peralta, Camille Wouters, Andrea L. Cox, Dorry L. Segev, Christine M. Durand, Mark Robien, Aaron A.R. Tobian, Andrew H. Karaba, Joel N. Blankson, William A. Werbel, Andrew Pekosz, Sabra L. Klein

**Affiliations:** W. Harry Feinstone Department of Molecular Microbiology and Immunology, Johns Hopkins University Bloomberg School of Public Health, Baltimore, Maryland; Department of Medicine, Johns Hopkins University School of Medicine, Baltimore, Maryland; Department of Surgery, New York University Grossman School of Medicine and NYU Langone Health, New York, NY; National Institute of Allergy and Infectious Diseases, Bethesda, MD; Department of Pathology, Johns Hopkins University School of Medicine, Baltimore, Maryland

## Abstract

Solid organ transplant recipients (SOTRs) suffer more frequent and more severe infections due to their compromised immune responses resulting from immunosuppressive treatments designed to prevent organ rejection. Pharmacological immunosuppression can adversely affect immune responses to vaccination. A cohort of kidney transplant recipients (KTRs) received their third dose of ancestral, monovalent COVID-19 vaccine in the context of a clinical trial and antibody responses to the vaccine strain, as well as to Omicron variants BA.1 and BA.5 were investigated and compared with healthy controls. Total IgG and live virus neutralizing antibody titers were reduced in KTRs compared to controls for all variants. KTRs displayed altered IgG subclass switching, with significantly lower IgG3 antibodies. Responses in KTRs were also very heterogeneous, with some individuals showing strong responses but a significant number showing no Omicron-specific neutralizing antibodies. Taken together, immune responses after COVID-19 vaccination in KTRs were not only lower than healthy controls but highly variable, indicating that simply increasing the number of vaccine doses alone may not be sufficient to provide greater protection in this population.

**Importance:** This study addresses the challenges faced by kidney transplant recipients (KTRs) in mounting effective immune responses against COVID-19. By evaluating the antibody responses to a third dose of monovalent mRNA COVID-19 vaccine and its effectiveness against Omicron subvariants (BA.1 and BA.5), this study reveals significant reductions in both binding and neutralizing antibodies in KTRs compared to healthy controls. The research highlights altered IgG subclass switching and heterogeneous responses within the KTR population. Reduced recognition of variants, coupled with differences in IgG subclasses, decreases both the quality and quantity of protective antibodies after vaccination in KTRs. These findings underscore the need for tailored vaccination strategies for immunosuppressed populations such as KTRs. Alternative formulations and doses of COVID-19 vaccines should be considered for people with severely compromised immune systems, as more frequent vaccinations may not significantly improve the response, especially regarding neutralizing antibodies.

## Introduction

Solid organ transplant recipients (SOTRs) have an elevated risk of severe COVID-19 infection and mortality due to immunosuppressive medications administered post-transplantation^1-5^. Following the initial two-dose mRNA vaccine series, many SOTRs exhibit weakened humoral and cellular immune responses against SARS-CoV-2^6-11^. Published studies on the third dose mRNA vaccine in SOTRs showed increased total anti-Spike (S) IgG antibodies and neutralizing antibodies compared to the initial dose series, suggesting higher immunogenicity^12-16^. While previous studies have extensively examined binding and neutralizing responses in this population, this study focuses on antibody quality-Omicron subvariant-specific antibody and IgG subclass responses-compared to healthy controls (HCs). An in-depth analysis of S-binding antibody subtypes and live virus neutralizing antibody responses in kidney transplant recipients (KTRs, n=81) and HCs (n=11) (**Supplementary Table S1**) was performed to understand KTR responses to the vaccine and antigenically distinct SARS-CoV-2 variants to determine the effectiveness of a third vaccine dose in this highly vulnerable population.

## Methods

The Methods are detailed in the Supplementary Material. This study used 81 samples from the COVID-19 Protection After Transplant (CPAT) pilot trial, assessing the third dose of COVID-19 mRNA vaccine in KTRs (**Supplementary Table 1**) ^17^. Serum samples were collected pre-vaccination, 30-, and 90-day post-vaccination. Enzyme-linked immunosorbent assays (ELISAs) measured S-specific IgG and IgG subtypes against the vaccine and Omicron variants and microneutralization titers determined. Antibody responses were compared pre- and post-vaccination across SARS-CoV-2 variants using one-way repeated measures ANOVA with GraphPad Prism 8 and Stata 15.

## Results

### KTRs mount lower vaccine-induced serological responses than healthy controls (HCs) after receipt of a third dose vaccine

Titers of antibodies to the S protein of the ancestral, Omicron BA.1 and Omicron BA.5 increased at 30 and 90 days post vaccination, though not all individuals seroconverted after receiving a third COVID vaccine (Figure 1). Live virus neutralizing antibody (nAb) titers also increased against ancestral and Omicron BA.1 and BA.5 in a portion of the population, but the number of non-responders was significantly greater in this functional assay when compared to total S protein binding antibodies (Figure 1B).

**Figure 1.**
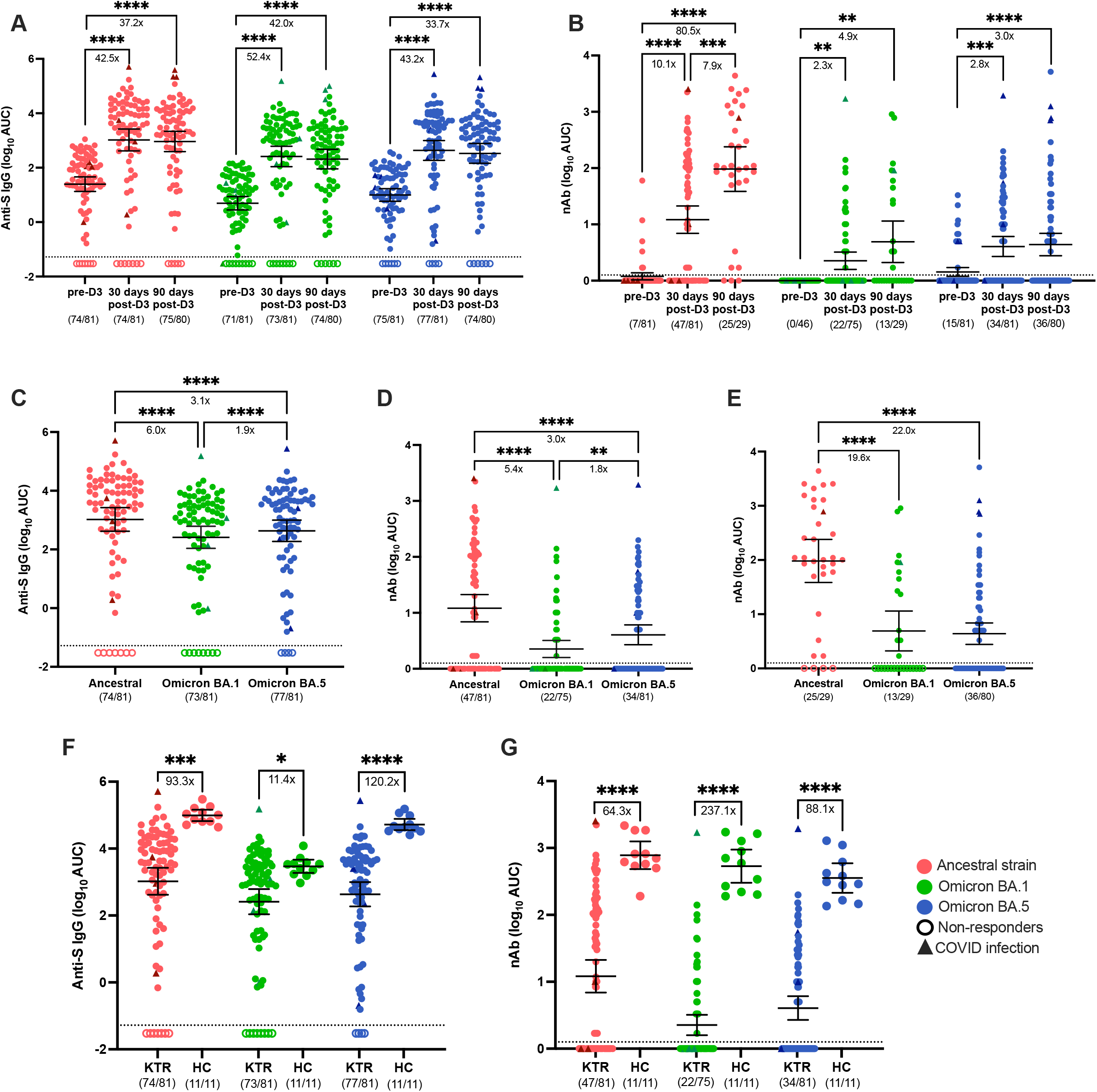
Serum IgG and nAb responses to SARS-CoV-2 ancestral and Omicron variants in KTRs in comparison of HCs. Total SARS-CoV-2 Spike (S)-specific IgG (**A**) and neutralizing antibody (nAb) (**B**) against the ancestral strain (red), Omicron BA.1 (green), and Omicron BA.5 (blue) variants measured in KTR (n=81) prior to dose 3 (pre-D3), 30 days post-dose 3 (30 days post-D3), and 90 days post-dose 3 (90 days post-D3). Total anti-S IgG (**C**) and nAb levels (**D**) at 30 days post-D3 and nAb levels at 90 days post -D3 (**E**) were compared between the ancestral strain, Omicron BA.1, and Omicron BA.5 variants. Total anti-S IgG (**F**) and nAb (**G**) against ancestral strain, Omicron BA.1, and Omicron BA.5 variants were compared between KTR and HCs (n=11) at 30 days post-D3. Dotted lines indicate the limit of detection (LOD), which is -1.52 for the anti-S IgG ELISA assay (**A, C, F**) and 0.17 for the nAb assay (**B, D, E, G**). Open circles represent non-responders with negative serological responses that fall below the LOD value. Solid triangles represent patients with a confirmed SARS-CoV-2 infection during the course of the study. The mean± 95% CI are shown in each panel. Significance is tested using one-way repeated measures ANOVA **(A-E**), and unpaired t-tests (**F, G**). *p < 0.05, **p < 0.01, ***p < 0.001, and ****p <0.0001. Fold changes (x) are labeled below the significance lines. Number of positive samples out of the total number of samples tested are indicated in parentheses.

When S protein antibody titers among the three SARS-CoV-2 variants were compared, the dose 3 (D3) responses to Omicron BA.1and Omicron BA.5 were significantly lower compared to the ancestral strain (**Figure 1C**). Similar results were observed when comparing the 90 days post-D3 data. The nAb responses to Omicron BA.1 and Omicron BA.5 remained significantly lower than the ancestral strain at 30 (Figure 1D) and 90 days (**Figure 1E**) post-D3, with a significant reduction in reactivity to Omicron BA.1 and Omicron BA.5 at 90 days post-D3. In our study, four participants with confirmed COVID-19 infection showed higher anti-S and nAb responses, but these outliers did not appear to significantly skew the results in this study population.

We next compared responses between vaccinated KTRs and HCs at 30 days post-D3. After receipt of the third mRNA vaccine, 100% of HCs and 90% of KTRs had detectable anti-S IgG responses against the three Spike variants (**Figure 1F**). However, KTRs had consistently lower antibody titers against ancestral S (93.3-fold decrease), BA.1 (11.4-fold decrease) and BA.5 (120.2-fold decrease) when compared to HCs (**Figure 1F**). All HCs had detectable nAb responses against all variants post-D3, but among KTRs, the nAb responders were only 47/81 to ancestral virus, 22/75 to BA.1 (), and 34/81 to BA.5 (**Figure 1G**). As compared with HCs, KTRs showed significant reductions in nAb titers against ancestral S (64.3-fold decrease), BA.1 (237.1-fold decrease), and BA.5 (88.1-fold decrease) (**Figure 1G**). Taken together, these data suggested that KTRs mounted improved antibody responses to a third dose of mRNA vaccine, but the titers remained lower and a lower proportion of KTRs mounted detectable responses.

### KTRs mount lower SARS-CoV-2 Spike specific IgG subclasses than healthy controls (HCs) after a third dose of vaccine

The abundance of subclass-specific IgG antibodies to SARS-CoV-2 ancestral S was assessed in KTRs after D3. There were differences in the proportion of individuals who generated subclass specific antibody responses post-D3. The IgG1, IgG3 and IgG4 responses were the strongest in this cohort, with IgG3 responses showing the lowest increase at days 30 and 90 post-D3 (**Figure 2A**). IgG1 and IgG3 subclass responses were strongest at 30 days post-D3 compared to IgG2 or IgG4 (**Figure 2B**). The abundance of IgG subclasses changed at 90 days post-D3 with IgG1 responses being the strongest, comparable IgG3 and IgG4 subclass levels and IgG2 responses continuing to be the lowest (**Figure 2C**). Overall, among KTRs, there was an increase in all IgG subclasses post-D3, with IgG1 as the dominant subclass in KTRs after D3 and IgG2 being the weakest response (**Figure 2D**).

**Figure 2.**
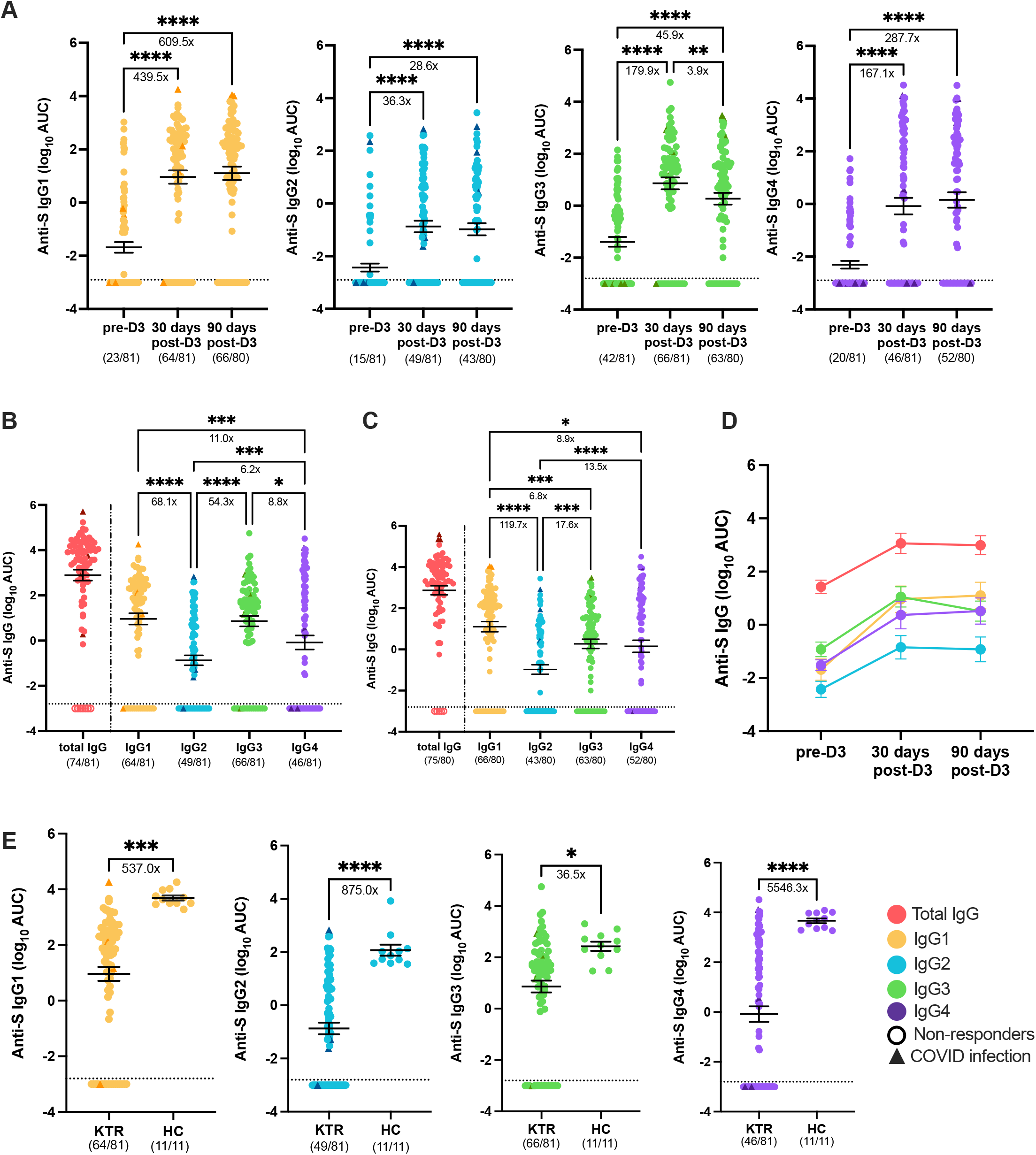
Serum IgG subclass profile to SARS-CoV-2 ancestral strain in KTRs. SARS-CoV-2 Spike (S)-specific IgG subclasses (IgG1-4): IgG1 (yellow), IgG2 (blue), IgG3 (green), and IgG4 (purple) against ancestral strain were measured in KTRs (n=81) prior to dose 3 (pre-D3), 30 days post-dose 3 (30 days post-D3), and 90 days post-dose 3 (90 days post-D3) (**A**). Anti-S IgG subclasses antibody levels against SARS-CoV-2 ancestral strain were compared at 30 days post-D3 (**B**) and 90 days post-D3 (**C**) in KTRs with anti-S total IgG (red) as a reference on the left of both panels. A summary panel of anti-S IgG subclass-specific antibody levels against ancestral strain are shown and connected by lines to show the changes of serological responses in IgG subclasses from pre-D3, 30 days post-D3, to 90 days post-D3 (**D**). Comparison of anti-S IgG subclass-specific antibody levels against ancestral strain were made between KTRs and healthy controls (HCs) (n=11) at 30 days post-D3 (**E**). Dotted lines indicate the limit of detection (LOD), which is -3.00 for the subclass-specific IgG ELISA assay. Open circles represent non-responders with negative serological responses that fall below the LOD value. Solid triangles represent patients with a confirmed SARS-CoV-2 infection during the course of the study. The mean± 95% CI are shown in each panel. Significance is tested using mixed-effects model (**A**), one-way repeated measures ANOVA (**B, C**), and unpaired t test (**E**). *p < 0.05, **p < 0.01, ***p < 0.001, and ****p <0.0001. Fold changes (x) are labeled below the significance lines. Number of positive samples out of the total number of samples tested are indicated in parentheses.

We next compared IgG subclass responses between KTRs and HCs 30 days post-D3 to determine if there were differences in vaccine-induced IgG subclasses between immunocompromised KTRs and immunocompetent individuals. KTRs had lower levels of all four IgG subclasses (**Figure 2E**), with IgG4 responses being particularly depressed. These findings suggested that KTRs mounted significantly lower anti-S IgG responses across all subclasses than HCs after a third dose of the COVID-19 mRNA vaccine.

## Discussion

The initial two doses and/or a third dose of a COVID-19 vaccine elicited anti-S IgG and nAb responses against the Omicron variants in many, but not all immunocompromised populations, with the level of serological responses usually lower when compared to healthy individuals. In KTRs, due to moderately to severely immunosuppressive status, a lower neutralizing activity and reduced S-specific IgG responses were determined when compared with HCs. These data demonstrate that the overall quantity and quality of COVID-19 vaccination and boosting induced antibody responses is not as great in KTRs compared to HCs and that differences still exist after a third vaccine dose. While nAb levels are one corelate of protection, the role of non-neutralizing antibodies and antibody Fc region functions in modulating COVID-19 disease severity remains poorly defined. Antibody Fc regions mediate activation of NK cells and macrophages in addition to fixing complement and mediating antibody dependent cellular cytotoxicity (ADCC) and enhancing the production of specific IgG subclasses may help improve protection from COVID-19 disease. Consideration should be given to different formulations and doses of COVID-19 vaccines in people with severely compromised immune systems, as more frequent vaccinations may not significantly increase the non-responding group, particularly when it comes to neutralizing antibodies.

## Acknowledgements

This work was supported by the Ben-Dov family, the Johns Hopkins COVID-19 Vaccine-related Research Fund, the National Cancer Institute (U54CA260492 (SLK), K24AI144954 (DLS), K08AI156021 (AHK), U01AI138897-05S (CMD/DLS), K23AI157893 (WAW), HHS N272201400007C (AP), HHS 75V593021C00045 (AP) and R01AI120938S1 (AART) from the National Institute of Allergy and Infectious Disease. We thank the participants for agreeing to be part of the study.

