## supplemental methods for "A Third COVID-19 Vaccine Dose in Kidney Transplant Recipients Induces Antibody Response to Vaccine and Omicron Variants but Shows Limited Ig Subclass Switching"

**Supplementary Table 1.** Baseline characteristics, vaccine platform and immunosuppressive regimen of kidney transplant recipient (KTR) and healthy control (HC)

|  | **Kidney Transplant Recipient**  **(KTR)** | **Healthy Control**  **(HC)** |  |
| --- | --- | --- | --- |
|  | Total (N=81) | Total (N=11) |  |
| **Demographics** |  |  | **p-value**† |
| **Age, no.(%)** |  |  | <0.001 |
| median (IQR) | 66 (57,73) | - |  |
| 18-49 | 13 (16) | 8 (73) |  |
| 50-64 | 23 (28) | 3 (27) |  |
| ≥65 | 45 (56) | 0 (0) |  |
| **Sex, no.(%)** |  |  | 0.777 |
| Female | 26 (32) | 4 (36) |  |
| Male | 55 (68) | 7 (64) |  |
| **Race, no.(%)** |  |  |  |
| White | 49 (60) | - | - |
| Black/African American | 24 (30) | - |  |
| Asian | 7 (9) | - |  |
| **Ethnicity, no.(%)** |  |  |  |
| Hispanic/Latino | 3 (4) | - | - |
| Non-Hispanic/Latino | 79 (96) | - |  |
| Prefer not to answer | 0 (0) | - |  |
| **COVID-19 and vaccination history** |  |  |  |
| **Prior SARS-CoV-2 infection*, no.(%)** | 4 (5) |  |  |
| **Days between 2nd and 3rd dose, median (IQR)** | 167 (149,177) |  |  |
| **Vaccine manufacture, no.(%)** |  |  |  |
| Pfizer-BioNTech (BNT162b2) | 59 (73) |  |  |
| Moderna (mRNA-1273) | 22 (27) |  |  |
| **Transplant history** |  |  |  |
| **Years since transplant, median (IQR)** | 5.4 (2.1,10.5) |  |  |
| **Medical comorbidities, no.(%)** |  |  |  |
| Diabetes | 26 (32) |  |  |
| HCV infection | 4 (5) |  |  |
| Lung disease | 16 (20) |  |  |
| Cardiovascular disease | 72 (89) |  |  |
| Autoimmune disease | 8 (10) |  |  |
| **Transplant immunosuppression** |  |  |  |
| **Baseline immunosuppressant, no. (%)** |  |  |  |
| Mycophenolate Mofetil | 55 (68) |  |  |
| Mycophenolic Acid | 9 (11) |  |  |
| High dose mycophenolate | 14 (22) |  |  |
| Prednisone | 75 (93) |  |  |
| Tacrolimus | 74 (91) |  |  |
| Cyclosporine | 5 (6) |  |  |
| **Triple IS, no. (%)**** | 57 (70) |  |  |
| * By positive prior molecular testing or reactive anti-nucleocapsid antibody at enrollment. ** Any combination of three transplant immunosuppressants (triple therapy) with a low-dose cyclosporine, azathioprine, and prednisone immunosuppression regimen at Day 0.  † Categorical variables (age category, gender, race, etc) were compared between SOTR cohort and control by chi-square tests  IQR, interquartile range; Triple IS, triple immunosuppression | | | |

**Methods**

**Study Background and design**

This study utilized samples from the COVID-19 Protection After Transplant (CPAT) pilot trial funded by the National Institutes of Health, a single-arm open-label trial of the safety and immunogenicity of a third dose of SARS-CoV-1 mRNA vaccination in kidney transplant recipients who had failed to respond to 2 prior mRNA vaccinations (NCT04969263)^1^. Serum samples were collected pre-vaccination, 30-days, and 90-days following the third dose COVID-19 mRNA vaccine (D3) (**Table 1**). Responses were compared with 11 healthcare workers (HC) who also received a third dose of mRNA vaccine, who served as immunocompetent controls (**Table 1**).

**Enzyme-linked immunosorbent assays (ELISAs)**

Standardized and validated indirect ELISAs were used to measure the titer of Spike (S)-specific IgG against the vaccine strain, and Omicron variants of SARS-CoV-2, as previously described ^2-6^. Recombinant SARS-CoV-2 S (2 ug/ml) engineered at Johns Hopkins or obtained through NCI Serological Sciences Network for COVID-19 were used to coat 96-well ELISA plates (Immulon 4HBX, Thermo Fisher Scientific) and were incubated overnight at 4 °C. Plates were washed with phosphate buffered saline with tween 20 (PBST) wash buffer (Thermo Fisher Scientific), blocked with 3% nonfat milk solution, and incubated for one hour at room temperature (RT). After incubation, the blocking buffer was discarded. Three-fold serially diluted serum samples, monoclonal antibody against the SARS- CoV-2 S protein (positive control; catalog 40150-D001, Sino Biological), and negative control samples were added and incubated for two hours at RT. Plates were washed with PBST, and horseradish peroxidase (HRP)- conjugated secondary IgG antibody (catalog A18823, Invitrogen, Thermo Fisher Scientific) was added at a 1:5,000 dilution and incubated for one hour at RT. For characterization of subclass-specific IgG, we used secondary IgG1 antibody (catalog 9054-05, SouthernBiotech) at a 1:4,000 dilution, IgG2 antibody (catalog 9060-05, SouthernBiotech) at a 1:4,000 dilution, IgG3 antibody (catalog 9210-05, SouthernBiotech) at a 1:4,000 dilution, and IgG4 antibody (catalog 9200-05, SouthernBiotech) at a 1:8,000 dilution. Plates were washed with PBST, and reactions were developed by adding SIGMAFAST OPD (o-phenylenediamine dihydrochloride) solution (MilliporeSigma), followed by a ten-minute incubation at RT. Reactions were stopped by adding 3M hydrochloric acid solution (Thermo Fisher Scientific). The optical density (OD) of each plate was read at 490 nm wavelength on a SpectraMax i3 ELISA Plate Reader (BioTek Instruments). Results were expressed as the log10-transformed area under the curve (AUC) generated from the background- subtracted OD values of the ten three-fold serial dilutions, as previously described ^5^. The limit of detection (LOD) was determined to be half of the lowest measured AUC with a corresponding endpoint titer of at least 20.

**Cells and Viruses**

Vero-E6-TMPRSS2 cells were obtained from the cell repository of the National Institute of Infectious Diseases, Japan and were grown in complete media (CM) consisting of Dulbecco's modified eagle medium (DMEM) containing 10% fetal bovine serum (FBS) (Gibco, Thermo Fisher Scientific), 1 mM glutamine (Invitrogen, Thermo Fisher Scientific), 1 mM sodium pyruvate (Invitrogen, Thermo Fisher Scientific), 100 U/mL penicillin (Invitrogen, Thermo Fisher Scientific), and 100 μg/mL streptomycin (Invitrogen, Thermo Fisher Scientific). Cells were incubated at 37°C in a humidified incubator with 5% CO2.

SARS-CoV-2 isolates B.1 (hCoV-19/USA/DC-HP00007/2020; B.1, GISAID EPI_ISL_434688), Omicron BA.1 variant (SCV2/USA/MD-HP20874/2021; BA.1.18, GISAID EPI_ISL_7160424), and Omicron BA.5 (hCoV-19/USA/MD-HP32103-PIDCNSQVGY/2022 GISAID EPI_ISL_15013106) were isolated from Vero-E6-TMPRSS2 cells plated in 24-well plates and grown to 75% confluence ^7^. The CM was removed and replaced with 150 μL of infection media (IM). IM is identical to CM except with an FBS concentration of 2.5%. 150 μL of the virus transport media containing a SARS-CoV-2 positive clinical swab was added to the culture. The cultures were incubated at 37°C for 2 hours, the inocula were aspirated and replaced with 500 μL of IM, and the cells were incubated at 37°C for up to 5 days. When cytopathic effect (CPE) was visible in most of the cells, the IM was collected and stored at –65°C. The presence of SARS-CoV-2 was verified by extracting RNA from the harvested supernatant using the QIAGEN QIAamp Viral RNA extraction kit and viral RNA was detected using quantitative rt-PCR. SARS-CoV-2 whole genome sequencing was performed on the cell culture virus isolates based on NGS technology. The consensus sequence of the virus isolate did not differ from the sequence derived from the clinical specimen. The infectious virus titer was determined on Vero-TMPRSS2 cells using a 50% tissue culture infectious dose (TCID50) assay ^8^. Serial 10-fold dilutions of the virus stock were made in IM, and 100 μL of each dilution was then added to the cells in a 96-well plate in hexaplicate. The cells were incubated at 37°C for 4-5 days, fixed with 4% formaldehyde for at least 4 hours and stained with naphthol blue-black overnight. Plates were scored visually for CPE. The TCID50 per mL was determined using the Reed and Muench calculation.

**Microneutralization assay**

Plasma nAbs were determined as described for SARS-CoV and modified for SARS-CoV-2 ^6, 9^. Two- fold dilutions of plasma, starting at a 1:20 dilution, were made using IM. Infectious virus was added to the plasma dilutions at a final concentration of 1 × 10^3 TCID50/mL or 100 TCID50 per 100 μL. The plasma-virus solution was incubated at room temperature for 1 hour, and 100 μL of each dilution was added to 1 well of a 96-well plate of VeroE6-TMPRSS2 cells in hextuplicate. The cells were incubated for 6 hours at 37°C with 5% CO2. The inocula were then replaced with fresh IM, and the cells were incubated at 37°C with 5% CO2 for 3-4 days until CPE is evident in the negative controls. The cells were fixed by the addition of 100 μL of 4% formaldehyde for at least 4 hours at room temperature, and then stained with naphthol blue-black overnight. The nAb titer was calculated as the highest serum dilution that eliminated the CPE in 50% of the wells (NT50) and the AUC was calculated using GraphPad Prism.

**Statistical analyses**

Statistical calculations were performed using GraphPad Prism 8 (GraphPad Software) and Stata 15 (StataCorp). Data are shown as mean± 95% confidence interval (CI) except where otherwise indicated. Demographics and clinical characteristics were evaluated with descriptive statistics. Comparison between antibody responses at pre- and post-vaccination and comparison of antibody responses among SARS-CoV-2 VOC were analyzed using one-way repeated measures ANOVA test. A p value less than 0.05 was considered significant.
